# Epistasis and an oligogenic architecture underlie the strength of a plant-herbivore interaction

**DOI:** 10.64898/2026.09.14.751214

**Authors:** Cintia Beatriz Pérez-López, Frédérique White, Nanouk Abonnenc, Sandrine L’Anglais-Landry, Érika Milot, Andrew Macdonald, Pierre-Étienne Jacques, Yasuhiro Sato, Matthew Barbour

## Abstract

1. Interaction strength between consumers and resources is a fundamental driver of eco-evolutionary dynamics, and its genetic architecture is key to predicting the pace of such dynamics. Our understanding of this architecture is limited for three reasons: (i) defense phenotypes do not necessarily translate to interaction strength; (ii) interaction strength is difficult to measure in ways tied to consumer-resource theory; and (iii) association studies typically prioritize detecting large-effect loci rather than characterizing whether interactions are polygenic (many small-effect genes) or oligogenic (few large-effect genes).
2. We address these gaps by examining how genomic variation in the plant *Arabidopsis thaliana* influences interactions with the aphid herbivore *Lipaphis pseudobrassicae* in a greenhouse experiment. We grew 390 accessions spanning the species’ natural range and measured aphid per-capita population growth as our interaction-strength metric. To test whether a previously identified plant defense gene (*AOP2*) acts additively or epistatically with other defense loci, we integrated our estimates of aphid population growth with public data on defense chemistry and gene expression. To determine whether the genetic architecture of interaction strength was polygenic or oligogenic, we used a Bayesian sparse linear mixed model that partitions these two components.
3. *AOP2* showed epistasis with two other defense genes (*MAM1* and *GS-OH*), indirectly modifying aphid population growth via the production of a specific defense compound (glucosinolate 2-hydroxy-3-butenyl). More broadly, interaction strength showed an oligogenic architecture: top single nucleotide polymorphisms (SNPs) occurred at low minor allele frequencies (< 3%) and grouped into six linkage disequilibrium blocks, including candidate genes outside canonical plant defense pathways. These six blocks explained 47% of the genetic variance in interaction strength, with only 3% additional variance attributable to polygenic effects.
4. Taken together, these results show that plant genomic variation modifies herbivore interaction strength through epistasis among defense genes and rare alleles at few loci. This oligogenic architecture imposes multiple constraints on the potential for rapid evolution, highlighting the need to integrate genomic effects into eco-evolutionary models.

## Introduction

Consumer-resource interactions are arguably the fundamental unit of ecological communities, as their strength and organization influence population dynamics, community resilience, and the flow of energy through ecosystems (McCann, 2012; Murdoch et al., 2003). The strength of a trophic interaction also plays a major role in determining the magnitude of natural selection (Benkman, 2013), thus shaping (co)evolutionary dynamics over space and time (Thompson, 2005). Such evolutionary change can feedback to modify interaction strength, which in turns alters the population dynamics of consumers and their resources (Becks et al., 2010; Cortez et al., 2020; Yoshida et al., 2003). Although there is accumulating evidence for the role of intraspecific genetic variation and rapid evolution in shaping consumer-resource interactions (i.e., evo-to-eco effects; Hendry, 2017), we still have a limited understanding of their genetic basis (Rudman et al., 2018). Yet, theory suggests that genetic architecture can play a key role in constraining or accelerating the pace of eco-evolutionary dynamics (Yamamichi, 2022), highlighting the importance of understanding their genetic underpinning.

Our current understanding of the genetic basis of trophic interactions is limited for three main reasons. First, although evolution is fundamentally a genetic process, much research on eco-evolutionary dynamics relies on the ‘phenotypic gambit’ (Grafen, 1984)— assuming that a trait’s underlying genetics can be ignored (Yamamichi, 2022). While often defensible (Rausher & Delph, 2015), this approach may fail when the trophic interaction is influenced by epistasis (gene-gene interactions) or a few loci of large effect. Indeed, well studied metabolic and putative defense traits are frequently determined by epistasis or large-effect loci (Barrett et al., 2008; Kliebenstein, 2017; Rowe & Kliebenstein, 2008; Steiner et al., 2007), suggesting that the phenotypic gambit can be misleading. It is also difficult to identify the specific traits mediating genetic effects on trophic interactions even after extensive phenotyping (Barbour et al., 2016), likely because researchers are biased by the tools available to measure traits. Trophic interactions may also be influenced by multiple traits (Barbour et al., 2015), which further complicates attempts to dissect their genetic architecture. Together, these factors suggest it may be more profitable to study the genetic basis of trophic interactions directly by treating the interaction itself as the phenotype of interest— an approach often used in studies of ‘community genetics’ (Bailey et al., 2006; Barbour & Pérez-López, 2025).

Second, eco-evolutionary dynamics between consumers and resources are fundamentally influenced by interaction strength, which is notoriously difficult to measure in a way that is strongly tied to ecological theory (Berlow et al., 2004; Wootton & Emmerson, 2005). At its core, measuring interaction strengths can be boiled down to quantifying how changes in the abundance (eco-to-eco effect) or phenotype (evo-to-eco effect) of one species alters the per-capita population growth of another (Berlow et al., 2004; Patel et al., 2018; St-Pierre et al., 2025). All other metrics of interaction strength with a clear theoretical basis can be derived from per-capita metrics of interaction strength (Berlow et al., 2004). While a growing number of genome-wide association studies (GWAS) use species’ abundances or community metrics as the phenotype of interest (Barbour & Pérez-López, 2025), these snap-shots from the field do not readily translate into per-capita metrics of interaction strength. Moreover, surveying a large genetic panel is difficult, which is why GWAS of specific trophic interactions often use proxies for interaction strength, such as survival or individual growth (Kloth et al., 2012), which does not always translate to per-capita population growth (McPeek & Peckarsky, 1998).

Finally, standard GWAS approaches rarely attempt to directly quantify genetic architecture (i.e., the number of loci and their distribution of effect sizes). For example, GWAS is often used to identify candidate loci of large effect but does not account for linkage disequilibrium among these loci (Zhou & Stephens, 2012). GWAS can give estimates of narrow-sense heritability (so called ‘SNP heritability’), but this metric necessarily assumes that many genes with small, additive effects are at play (Speed et al., 2017; Stanton-Geddes et al., 2013). Yet, the genetic architecture of trophic interactions may lie on a continuum between ‘many genes of small effect’ and ‘few genes of large effect’ (Zhou et al., 2013), as expected for local adaptation (Orr, 2005). Standard GWAS also assumes that all loci have additive effects (Zhou & Stephens, 2012). However, as noted above, well-studied metabolic and defense traits are often governed by epistasis among genes (Kliebenstein, 2017; Rowe et al., 2008; Steiner et al., 2007), which standard GWAS is not adapted to detect. Taken together, we need research that characterizes the architecture of genetic effects on the per-capita population growth rates of interacting species. Doing so will yield fundamental insight into how genetic architecture dictates the pace and potential for eco-evolutionary dynamics to unfold and allow the application of existing theory down to the level of individual genes.

Here, we sought to characterize the genetic architecture underlying the strength of a plant-herbivore interaction. To do this, we measured the per-capita population growth of the aphid herbivore *Lipaphis pseudobrassicae* across 390 accessions of the plant *Arabidopsis thaliana*—capturing broad geographic and genomic variation (Fig. 1)— in a greenhouse experiment. Previous work showed that allelic variation at the plant defense gene *AOP2* altered multi-trophic food-web persistence through its effects on *L. pseudobrassicae* population dynamics (Barbour et al., 2022). *AOP2* is a key player in the production of aliphatic glucosinolates, a diverse class of secondary metabolites involved in insect and pathogen defense in the Brassicaceae (Kliebenstein, 2017; Kliebenstein et al., 2001). Specifically, the presence of a functional allele at *AOP2* modifies the side chain of the core methylsulfinyl compounds to catalyze alkenyl glucosinolate production (notably Allyl, 3-butenyl, and 2-OH-3-butenyl; Kliebenstein et al., 2001). However, the role of natural variation at *AOP2* across diverse genetic backgrounds is unclear, as prior work evaluated mutant lines of *A. thaliana* in a single genetic background (Barbour et al., 2022). Moreover, *AOP2* interacts epistatically with structural variation at other glucosinolate genes (e.g., *MAM1* and *GS-OH*) to produce specific compounds (Hansen et al., 2008; Kliebenstein, Kroymann, et al., 2001; Kliebenstein & Cacho, 2016; Kroymann et al., 2003), but whether the effects of *AOP2* on *L. pseudobrassicae* are additive or epistatic with other glucosinolate genes is unknown. Finally, prior work highlighting the importance of *AOP2* only tested its effect relative to a few plant defense genes (Barbour et al., 2022); therefore, whether natural variation across the genome is important in shaping the strength of this plant-herbivore interaction remains unknown.

**Figure 1.**
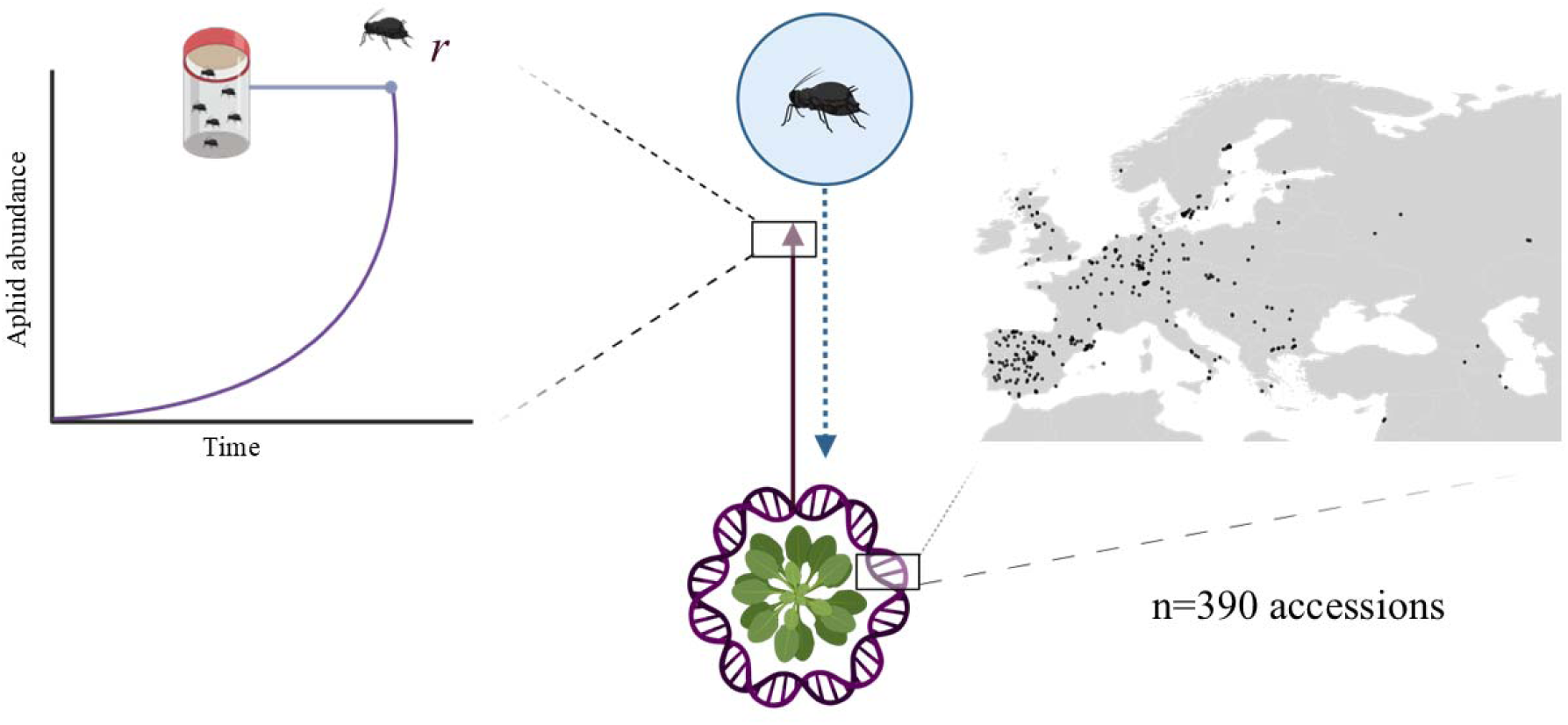
Study system and quantifying the genetic architecture of a plant-herbivore interaction. On each of 390 *Arabidopsis thaliana* accessions (representing broad geographic and genomic variation), we measured the per-capita population growth (*r*) of the aphid herbivore *Lipaphis pseudobrassicae*. Map indicates the geographic origin of each accession panel.

Using this study system, we sought to address two questions: (1) Does *AOP2* act additively or epistatically with other genes to affect aphid per-capita population growth? (2) What is the broader genetic architecture underlying the strength of this plant-herbivore interaction? To address the first question, we combined an *AOP2-*focused association study with publicly available data on glucosinolate gene expression and chemical concentrations for our diverse panel of accessions (Katz et al., 2021; Kawakatsu et al., 2016). If *AOP2* has additive effects on aphid population growth, we would expect the total production of *AOP2*-derived alkenyl glucosinolates (Allyl + 3-butenyl + 2-OH-3-butenyl) to be the primary driver. Conversely, if *AOP2* effects are epistatic to other glucosinolate genes, we would expect specific alkenyl glucosinolates (Allyl vs. 3-butenyl vs. 2-OH-3-butenyl) to better predict aphid population growth. To address the second question, we used a Bayesian sparse linear mixed model (BSLMM) to quantify the number of loci and the relative contribution of large- and small-effect genes, while simultaneously accounting for linkage disequilibrium. Taken together, this work leverages an experimentally tractable system to identify the genetic basis of plant-herbivore interaction strength.

## Materials and methods

### Plant genotypes

To characterize the broad range of natural genomic variation in *A. thaliana*, we initially selected 512 accessions from the 1001 Genomes Project (Alonso-Blanco et al., 2016) that had been filtered from the initial pool of 1,135 available accessions to ensure high-quality and genetically diverse samples (Exposito-Alonso et al., 2019). Specifically, this subset of accessions excluded those with low genome coverage (less than 10x), nearly identical genomes, and accessions from overrepresented geographic regions to minimize geographic bias, resulting in a more representative dataset of *A. thaliana*’s genomic variation (details given in Exposito-Alonso et al., 2019). Seeds were obtained from the Arabidopsis Biological Resource Center (https://abrc.osu.edu/) and grown for one generation to minimize maternal effects and have a sufficient number of seeds for the experiment (details on the growing conditions for this generation of plants are provided in Supporting Information: Bulking seeds). Of these 512 accessions, we selected 390 to provide a manageable experimental design. This number of accessions (>290) can achieve sufficient statistical power for detecting genes with large phenotypic effects (Fujii et al., 2019; Groux et al., 2021). These 390 accessions were chosen to maximize overlap with existing data from the AraPheno database (Seren et al., 2017) on gene expression (Kawakatsu et al., 2016), and defense chemistry (Katz et al., 2021), allowing us to explore the mechanistic links between genomic variation and herbivore interaction strength.

### Greenhouse experiment

To determine how plant genomic variation influenced plant-herbivore interactions, we conducted a greenhouse experiment with the previously described *A. thaliana* accessions. To ensure experimental feasibility and replication, we divided the experiment into three temporal blocks. Each block contained one replicate plant per accession, totalling 1,170 plants. Seeds were sown in 7 cm diameter pots filled with Pro-Mix HP Mycorrhizae Potting soil and stratified in the dark at 4°C for two weeks to promote simultaneous germination across these diverse accessions. Plants were then grown in the greenhouse under a cycle of 16 hours light at 22°C and 8 hours dark at 20°C for two weeks before adding two first-instar aphid nymphs to each plant using a fine paintbrush (details on aphid growing conditions before adding to the experiment provided in Supporting Information: Bulking aphids). To prevent aphid dispersal, we secured a fine-mesh organza bag (10 x 15 cm) with a rubber band over each potted plant. Because *L. pseudobrassicae* develop from first-instar nymphs to adults in approximately 7 days, we collected aphids after two weeks to measure their population growth. Aphids from each plant were collected in plastic bags, preserved at -20°C, and counted under a dissecting scope. To account for potential effects of plant size on plant-herbivore interactions (e.g., aphids may thrive better on larger plants), we measured the length (cm) of the cotyledons and true leaves one day before introducing aphids. Total plant area was estimated as an ellipse (Fig. S1), where the longest radius of the ellipse corresponds to half the cotyledon length, and the shortest radius corresponds to half the true leaf length: Area=π×(Cotyledons/2) × (True leaves/2). A constant of 0.1 was added to each measured length (cotyledon and leaf) before calculating the area to ensure positive values for seedlings where true leaves had not yet emerged. During this experiment, six of the 390 accessions experienced complete plant mortality, preventing reliable aphid count data from being collected. These accessions were therefore excluded, resulting in a final set of 384 accessions for which robust phenotypic values could be obtained (Table 1).

**Table 1.**
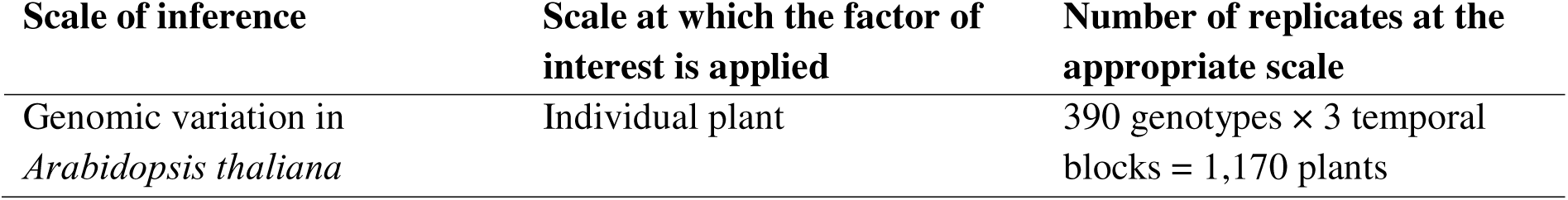
Replication statement.

| Scale of inference | Scale at which the factor of interest is applied | Number of replicates at the appropriate scale |
| --- | --- | --- |
| Genomic variation in *Arabidopsis thaliana* | Individual plant | 390 genotypes × 3 temporal blocks = 1,170 plants |

### Quantifying plant-herbivore interaction strength

Ideally, the interaction strength of an evo-to-eco effect should measure how the change in a heritable phenotype or allele (evolution) of one species affects the per-capita population growth (ecology) of the interacting species (Berlow et al., 2004; Patel et al., 2018; St-Pierre et al., 2025). We quantified aphid per-capita population growth (*r*) as the natural log difference between the final number of aphids on each plant () and the initial number of nymphs (): ln ln. First-instar nymphs are delicate and occasionally died after being transferred to experimental plants (see Supporting Information: Follow-up experiment to assess aphid mortality for details). To account for this source of mortality, we fit a Bayesian hierarchical model with a custom Poisson-mixture distribution to the raw aphid count data (). This custom distribution models the data as a mixture of a Poisson count process and a binary mortality process, assuming the two added nymphs die at rate *m*, allowing for varying probabilities of the initial population (i.e., 2, 1, or 0 nymphs). Plant accession ID and experimental block were modelled as fixed effects to avoid the double-shrinkage problem (“BLUPing twice”) that can bias subsequent genome-wide association studies (GWAS; Holland & Piepho, 2024). To account for overdispersion, we included an observation-level random effect (Harrison, 2014). The model was fit using the R package *brms* (Bürkner, 2017) with four Markov Chain Monte Carlo (MCMC) chains, each run for 6000 iterations, following a warm-up period of 3000 iterations. We specified weakly, regularizing priors (normal (mean = 0, SD = 2)) for fixed effects and default priors for random effect. Model convergence was assessed by ensuring that R-hat was below 1.01 for all parameters. We used the posterior mean for each accession as the Best Linear Unbiased Estimate (BLUE) of the herbivore’s per-capita population growth rate.

### Does AOP2 act additively or epistatically with other genes to affect aphid per-capita population growth?

To answer this question, we integrated our accession-specific estimates of aphid per-capita population growth with publicly available data on glucosinolate defense chemistry. Aliphatic glucosinolate concentrations in seeds from 797 European accessions grown under controlled conditions were sourced from (Katz et al., 2021). Compound concentrations were converted to mol/g to improve interpretability of model coefficients. To test whether aphid population growth was better explained by an additive or epistatic model, we used Aikaike Information Criterion (AIC) to compare two alternative linear regression models. The additive model fit the total concentration of *AOP2*-derived compounds (Allyl + 3-butenyl + 2-OH-3-butenyl) as the focal predictor, while the epistatic model included the three compounds separately as predictors. The epistatic model was simplified by removing compounds with non-significant effects (*p* > 0.05), and this final model was used for comparison to the additive model. All models included the average estimated plant size for each accession as a covariate to isolate the effects of chemical variation.

To further explore the mechanistic effects of *AOP2*, we modelled the direct and indirect effects of variation at glucosinolate genes, chemistry, and plant size on aphid population growth by fitting a structural equation model (SEM) with the R package *piecewiseSEM* (Lefcheck, 2015). To do this, we first identified top scoring SNPs at glucosinolate genes (SNP windows for each gene followed Katz *et al*. 2021) based on the GWAS (described in the next section) using the following criteria: uncorrected GWAS *p*-value < 0.05 and minor allele frequency > 0.1. We filtered based on minor allele frequency > 0.1 to facilitate testing of epistasis among glucosinolate genes. From these candidate SNPs, we retained the SNP that had the largest and clearest effect on its respective glucosinolate gene expression (gene expression data were sourced from the AraPheno database (https://arapheno.1001genomes.org; Seren et al., 2017). We then fit the following three linear models: 1) the effect of the top scoring SNPs on glucosinolate concentrations; 2) the effect of glucosinolate concentrations on plant size; and 3) the simultaneous effects of both glucosinolate concentrations and plant size on aphid per-capita population growth. All predictors and response variables for the SEM were standardized (mean = 0, SD = 1) to more easily compare model coefficients (denoted in Results section). Model adequacy was assessed using Fisher’s *C* tested on a chi-squared distribution with 2*k* degrees of freedom, where *k* is the number of missing paths in the directed separation test (Shipley, 2000). Note that *p* > 0.05 indicates an adequate SEM fit (i.e., no statistically significant missing paths).

### What is the broader genetic architecture underlying the strength of this plant-herbivore interaction?

To answer this question, we fit a Bayesian sparse linear mixed model (BSLMM) with the software GEMMA (Zhou & Stephens, 2012) to our accession-specific estimates of aphid population growth. BSLMM simultaneously models the variance explained by many genes with small, normally distributed effects (the polygenic component) and a smaller subset of genes with potentially larger, sparse effects (the sparse component), while accounting for population structure via a genetic relationship matrix (Zhou et al., 2013). We used imputed genotypes for 10,709,466 SNPs across 2,029 *A. thaliana* accessions that were obtained from the AraGWAS Catalog (https://aragwas.1001genomes.org/#/download-center; Togninalli et al., 2018). For analysis, we retained SNPs with a minor allele frequency > 0.01, resulting in 3,740,069 high-quality SNPs. We accounted for population structure by computing a centered genetic relationship matrix (“-gk 1” option) among accessions. We ran the BSLMM using the standard linear model (“-bslmm 1” option), which estimates SNP effect sizes and variance components using MCMC. We then ran three independent chains of the model with 2,500,000 burn-in steps, followed by 10,000,000 sampling steps where parameter values were recorded every 100 steps. The BSLMM estimates four key parameters to describe the underlying genetic architecture: the proportion of variance explained by all SNPs (PVE); the proportion of genetic variance explained by sparse effects (PGE); the number of SNPs with sparse effects (); and the SNP-specific Posterior Inclusion Probability (PIP), which quantifies the probability that an individual SNP has a non-zero effect (Zhou et al., 2013). We report estimates of model parameters aggregated across the three independent chains in the main text but report the results for each independent chain in Table S1.

To map SNPs to candidate genes, we examined local linkage disequilibrium (LD) around the top-associated SNPs identified by our BSLMM (PIP > 0.01). Pairwise linkage disequilibrium was calculated as the squared Pearson correlation coefficient () between biallelic markers, which ranges from 0 to 1 and measures the strength of association between two loci (Hill & Robertson, 1968; VanLiere & Rosenberg, 2008). We then performed hierarchical clustering using LD distance (1 - *r*²) as the dissimilarity measure, where values near 0 indicate strong LD and values near 1 indicate weak LD. The LD blocks were defined at a standard cut-off of ≥ 0.8 (Daly et al., 2001; Dong et al., 2021). To identify candidate genes, we then cross-referenced these coordinates with the Ensembl Plants genome browser (v.62), retrieving gene annotations for each LD block (Kinsella et al., 2011). Genes located within a LD block were considered candidate genes. When the signal did not overlap genes (i.e., intergenic region), the nearest genes upstream and downstream within a 10 kb search window (standard distance for capturing cis-regulatory targets in *A. thaliana*: (Kim & Jander, 2007; Nordborg et al., 2002), were retained as the primary candidates. This approach is consistent with common GWAS interpretation practice in cases where causal genes cannot be assigned directly (Buniello et al., 2019; Nasser et al., 2021; Uffelmann et al., 2021). To assess the robustness of the BSLMM results, we also conducted a genome-wide association study (GWAS) in GEMMA using linear mixed models. These analyses used the same filtered genotype dataset and centered genetic relationship matrix described above. SNP-trait associations were tested using the likelihood ratio test (“-lmm 2” option). We fit two GWAS models: one model without plant size as a covariate, to align with the BSLMM analysis, and a second model including plant size as a covariate to evaluate whether associations were robust to size-dependent effects.

All code was executed with R (version 4.4.0, R Core Team, 2025, http://www.r-project.org) or Python 3.11.5. Computational analyses for the BSLMM and GWAS were performed on the Digital Research Alliance of Canada high-performance computing infrastructure.

## Results

### Does AOP2 act additively or epistatically with other genes to affect aphid per-capita population growth?

Our additive model showed that the total concentration of *AOP2*-derived chemicals negatively affected herbivore population growth (coef. [95% CI] = -0.04 [-0.08, -0.01], *p* = 0.021). However, the model for individual chemicals revealed that this effect was driven primarily by 2-OH-3-butenyl (Fig. 2A). Specifically, herbivore population growth decreased by 16% across the natural range of variation in 2-OH-3-butenyl concentration (coef. [95% CI] = -0.06 [-0.10, -0.02], *p* =0.007). In contrast, the other *AOP2*-derived compounds did not clearly affect herbivore population growth (Allyl, coef. [95% CI] = 0.003 [-0.061, 0.067], *p* = 0.925; 3-butenyl, coef. [95% CI] = 0.02 [-0.16, 0.19], *p* = 0.860). Akaike’s Information Criterion (AIC) indicated that the epistatic model containing only the specific compound 2-OH-3-butenyl (AIC = 526.89) was a better predictor of aphid per-capita population growth than the additive model (AIC = 530.78).

**Figure 2.**
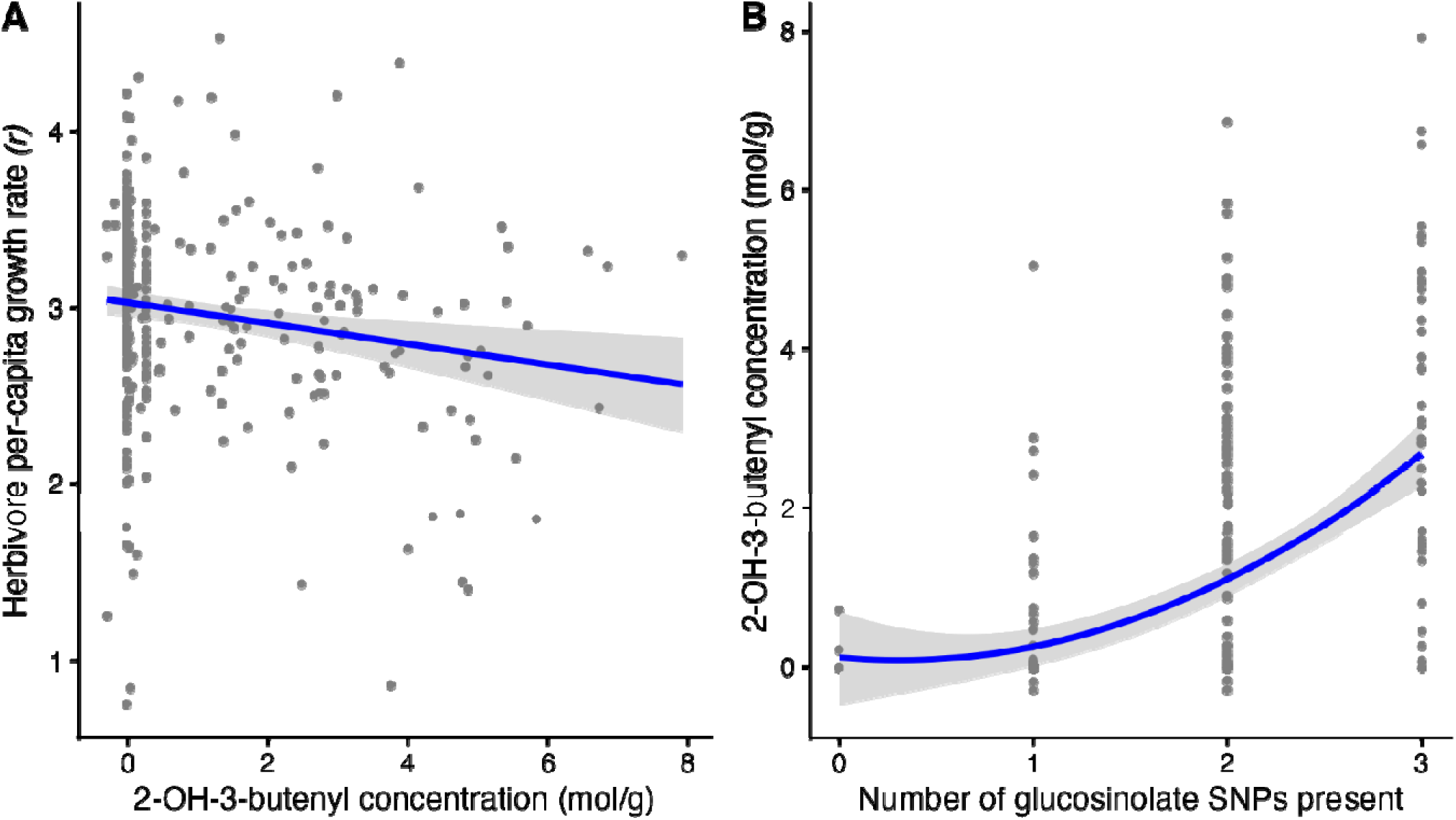
Epistasis in plant defense genes indirectly influence herbivore per-capita growth rate by altering the concentration of a specific chemical compound. (**A**) The concentration of the glucosinolate compound 2-OH-3-butenyl negatively affects herbivore per-capita population growth rate. (**B**) The number of glucosinolate SNPs present has non-additive effects on 2-OH-3-butenyl concentration. Each point corresponds to an accession of *Arabidopsis thaliana* (n=310). Solid lines and bands correspond to the mean and 95% confidence interval of each relationship.

Our structural equation model (SEM) revealed that epistasis among glucosinolate genes *AOP2*, *MAM1* and *GS-OH* strongly predicted the concentration of 2-OH-3-butenyl (Fig. 2B). Specifically, there was a clear quadratic effect of the number of glucosinolate SNPs on 2-OH-3-butenyl concentration (β [95% CI] = 0.90 [0.38, 1.42], *p* < 0.001), indicating that non-additive genetic effects explained 23% of the variance. Contrary to expectations of a growth-defense trade-off, 2-OH-3-butenyl concentration did not clearly affect plant size (β [95% CI] = 0.02 [-0.09, 0.13], *p* = 0.727). As predicted, herbivore population growth was jointly influenced by plant size and defense chemistry. Herbivore population growth increased with larger plants (β [95% CI] = 0.48 [0.38, 0.58], *p* < 0.001) and decreased with higher 2-OH-3-butenyl concentration (β [95% CI] = -0.15 [-0.25, -0.05], *p* = 0.002). Together, plant size and glucosinolate concentration explained 25% of the variance in herbivore population growth. Note, however, that the global goodness-of-fit test for our SEM was statistically significant (Fisher’s C = 16.77, df = 8, *p* = 0.011), indicating that the hypothesized model structure was missing one or more paths. Follow-up analyses indicated that there was an unexplained effect of additive SNP variation on aphid per-capita growth (β [95% CI] = -0.17 [-0.28, -0.06], *p* = 0.002) independent of 2-OH-3-butenyl concentration and plant size, suggesting that these glucosinolate SNPs may have additional negative effects on aphid population growth through a separate mechanism.

### What is the broader genetic architecture underlying the strength of this plant-herbivore interaction?

The Bayesian sparse linear mixed model (BSLMM) revealed that the strength of plant-herbivore interactions is a highly heritable trait, with host genetics explaining approximately half of the variance (median PVE [95% CI] = 0.50 [0.25, 0.78]). Moreover, the architecture was oligogenic, with a small number of large effect SNPs (median [95% CI] = 4 [2, 13]) predicted to explain most of the heritable variation (Fig. 3A; median PGE [95%CI] = 0.47 [0.21, 0.87]). Our model identified 33 SNPs with posterior inclusion probabilities (PIP) > 0.01, all of which were rare (minor allele frequencies < 0.03; Table S2). These top SNPs were distributed across six independent linkage disequilibrium blocks on chromosomes 1, 4, and 5 (Fig. 3A, Tables S3). This block structure aligns well with the estimated number of SNPs with sparse effects (), further supporting an oligogenic architecture (Fig. 3A). These linkage disequilibrium blocks were also reflected in a classic GWAS, with at least one SNP in 5 of 6 blocks passing the false-discovery rate threshold (green line in Fig. 3B) and 2 of 6 blocks passing a more stringent Bonferroni cutoff (pink line in Fig. 3B). In a GWAS that additionally controls for average plant size, only 3 of 6 blocks were above the false-discovery rate threshold (1 of 6 above Bonferroni cutoff), suggesting that the effect of some of these SNPs was at least partially mediated by their effect on plant growth (Fig. S2).

**Figure 3.**
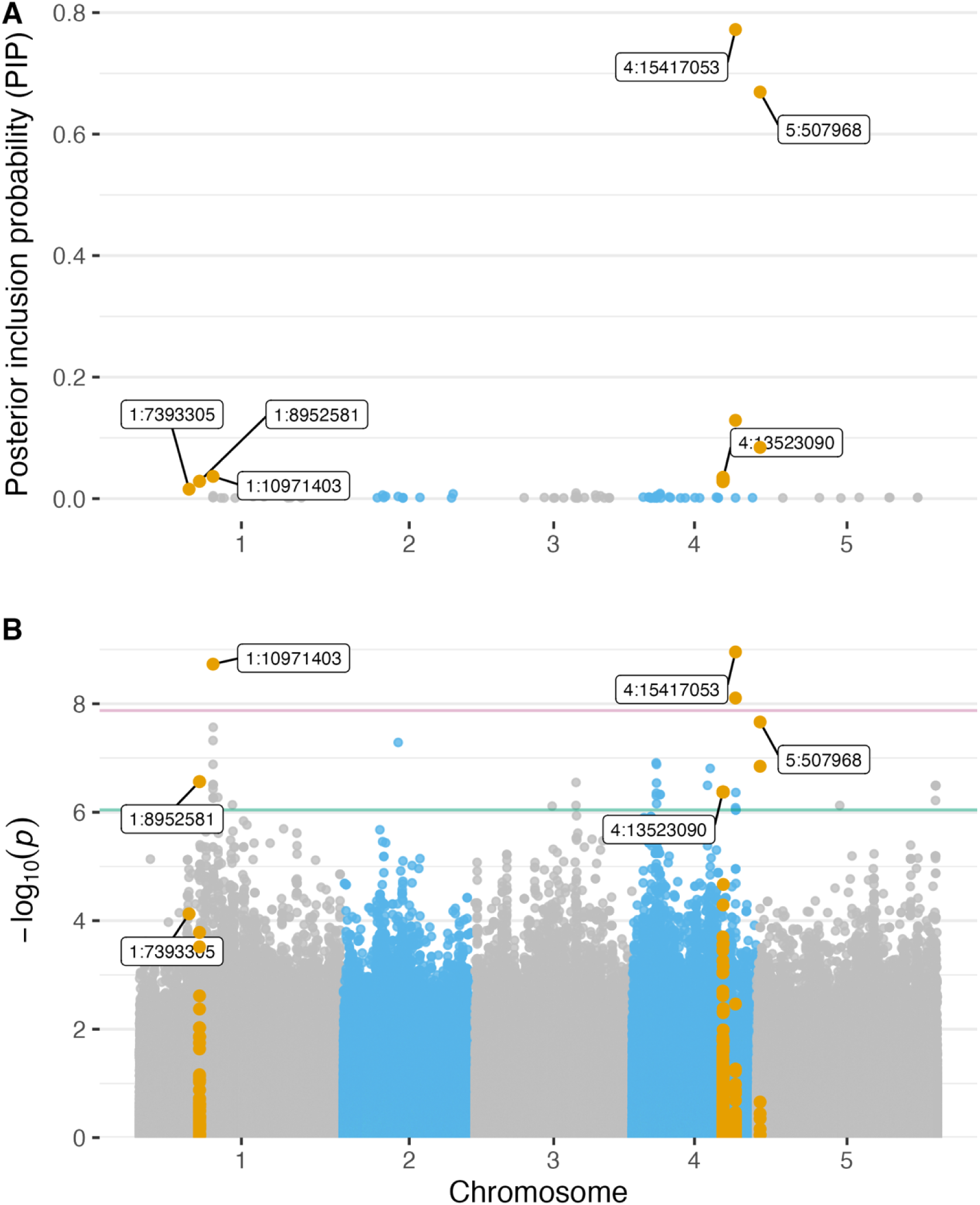
Genome-wide association results for plant-herbivore interaction strength. Manhattan plots of single-nucleotide polymorphism (SNP) associations in *Arabidopsis thaliana* with herbivore per-capita population growth from a Bayesian sparse linear mixed model (**A**) and a classic GWAS (**B**). Each point represents a SNP, with orange points highlighting SNPs with posterior inclusion probabilities (PIP) > 0.01 (i.e., probability of having a non-zero effect), as inferred from the BSLMM. Positions of SNPs with the lowest GWAS *p*-value in each of the six linkage disequilibrium blocks identified by BSLMM are labelled. The green and pink horizontal lines in (**A**) denote the genome-wide false-discovery rate and Bonferroni thresholds, respectively.

The six independent LD blocks varied markedly in structure. Specifically, blocks 2 and 4 overlapped with annotated genes, whereas blocks 1, 3, 5 and 6 were located in intergenic regions. The two SNPs with the highest probability of being retained in the model were in block 5 on chromosome 4 (4:15417053, PIP = 77.22%) and block 6 on chromosome 5 (5:507968, PIP= 66.94%; Figure 3A). Both of these SNPs were intergenic, suggesting they may tag regulatory variants. For block 5 on chromosome 4, the flanking region includes *GPX7* (glutathione peroxidase 7; Chang et al., 2009) and *PDS5* (cohesin complex component; Pradillo et al., 2015) (Table S3). For block 6 on chromosome 5, the flanking regions include *MT2B* (metallothionein 2B; Guo, Meetam, et al., 2008) and *DAU1* (DNA-damage induced 1; Borg et al., 2011) (Table S3). In contrast, the LD interval on block 2 of chromosome 1 contained three annotated genes: *RCN1* (ARM-repeat superfamily protein; Kwak et al., 2002), *AT1G25500* (plasma-membrane choline transporter family protein; Taylor et al., 2021), and *AT1G25510* (eukaryotic aspartyl protease family protein; Theologis et al., 2000). Block 4 on chromosome 4 was the largest block, spanning 76.3 kb that included 22 genes, none of which have been previously implicated in plant defense (Table S4).

## Discussion

Our results show that plant genetic architecture has a strong and structured effect on the strength of a plant-herbivore interaction. This structure was expressed through epistatic effects among three plant defense genes as well as additive effects concentrated in a limited number of loci that have not been previously implicated in plant defense. The epistatic interactions among the plant defense genes channel metabolic flux toward chemical specificity (Hansen et al., 2008; Kliebenstein, Kroymann, et al., 2001; Kliebenstein & Cacho, 2016; Kroymann et al., 2003), where the accumulation of a single compound (2-OH-3-butenyl) matters more than total pathway output. This pattern is consistent with the view that plant defense systems are organized as integrated molecular networks that scale to organismal and population-level outcomes (Kliebenstein, 2014). In addition, our genome-wide analysis identified an oligogenic architecture, in which a limited number of loci with additive effects explain approximately 47% of heritable variation in herbivore’s response. The mechanisms at these major-effect loci remain unknown, but their large effect sizes imply that they too may act through discrete, potent ecological mechanisms rather than diffuse quantitative variation. Together, these findings suggest that plant defense is not a diffuse background of many small additive effects (Hendry, 2013), but rather a system in which conditional pathway interactions and a small set of major genomic regions shape herbivore population growth.

### Does AOP2 act additively or epistatically with other genes to affect aphid per-capita population growth?

We found that *AOP2* exhibited epistasis with two other glucosinolate genes (*MAM1* and *GS-OH*) to indirectly modify aphid population growth through their effect on the production of a specific glucosinolate compound, 2-hydroxy-3-butenyl. Prior work with *Lipaphis* aphids has found that total glucosinolate concentrations have a negative effect on aphid population growth (Kumar et al., 2017), for which 2-hydroxy-3-butenyl is often a dominant compound (Katz et al., 2021). *Lipaphis* aphids are crucifer specialists that have evolved the ability to sequester glucosinolates, effectively turning into a ‘mustard oil bomb’ when they are attacked by generalist predators (Bridges et al., 2002). The related crucifer specialist, *Brevicoryne brassicae*, selectively avoids uptake of 2-hydroxy-3-butenyl (Goodey et al., 2015), suggesting that this glucosinolate compound may be particularly toxic. It’s worth noting that gene expression at *AOP3*, which is co-localized with *AOP2*, has been shown to be upregulated in response to herbivory from *Lipaphis* aphids (Sato et al., 2019); however, we did not find any evidence that *AOP3*-derived compounds negatively affected *L. pseudobrassicae* population growth (Table S4). Future work is needed to identify the molecular mechanisms underlying *Lipaphis pseudobrassicae*’s observed sensitivity to 2-hydroxy-3-butenyl.

Epistatic interactions among biosynthetic and regulatory loci have been documented as important determinants of chemical-defense phenotypes and metabolism across diverse plant systems (Hughes, 1991; Kliebenstein, 2017; Rowe & Kliebenstein, 2008; Zangerl & Berenbaum, 2004) and, more broadly, in affecting life-history traits of plants and animals (Burch et al., 2024). Secondary metabolite pathways are often organized as multistep biosynthetic cascades containing branch points, enzymes competing for shared substrates, and, in some cases, separate activation systems in which a toxin precursor and its hydrolytic enzyme are stored separately (Hughes, 1991; Kliebenstein, 2017). This structure creates natural conditions for epistasis because the phenotypic effect of variation at one locus, such as a branch-point enzyme, can depend on the genotype at another locus, such as an upstream enzyme supplying its substrate or a downstream enzyme activating its product. Epistasis may therefore be a recurring feature of the genetic architecture underlying chemical defense s, such that the phenotypic and ecological consequences of variation at one locus are frequently contingent on genotype at another. This genetic architecture could maintain genetic variation at chemical-defense loci by masking allelic effects from selection in some genetic backgrounds, but also constrain rapid evolution when the selective advantage of an allele depends on the presence of particular alleles at other loci (Barrett & Schluter, 2008). Consequently, standard quantitative-genetic models that predict evolutionary responses primarily from additive genetic variance, such as the breeder’s equation and the multivariate response-to-selection equation (Lande & Arnold, 1983), may provide incomplete predictions when selection acts on epistatically determined chemical-defense phenotypes.

### What is the broader genetic architecture underlying the strength of this plant-herbivore interaction?

Our genome-wide association analysis revealed that this plant-herbivore interaction is not a polygenic trait. Instead, it is governed by an oligogenic architecture, with rare alleles at a small number of loci explaining a large fraction of the additive genetic variance. Our LD block analysis of the top SNPs (PIP > 0.01) reduced these signals to six independent loci, with the strongest associations on chromosomes 4 and 5. While these SNPs reside in intergenic regions, making it difficult to pinpoint the causal genes without further fine-mapping, the nearest upstream and downstream genes provide candidates for future investigation. In GWAS, most trait-associated SNPs often fall outside coding regions, which may affect gene regulation, pointing to the functional roles of long non-coding RNAs (Wang et al., 2015). On chromosome 4, the associated region is flanked by *PDS5* and *GPX7*; on chromosome 5, the strongest region lies near *DAU1* and *MT2B*. None of these genes has a direct annotated role in chemical defense, but that is itself informative. *PDS5* is involved in chromosome cohesion and stress-related processes (Göbel et al., 2024; Pradillo et al., 2015), *GPX7* contributes to oxidative stress responses (Chang et al., 2009; Galant et al., 2011), *DAU1* has roles in DNA damage repair (Borg et al., 2011; Pradillo et al., 2015), and *MT2B* is involved in metal homeostasis and detoxification (Guo et al., 2003; Jie Lei et al., 2021). Their proximity to associated variants suggests that plant-herbivore interactions may be influenced not only by defense chemistry itself, but also by broader physiological processes such as stress responses, resource allocation, and plant vigour, which are all known to affect herbivore performance (Davila Olivas et al., 2017; Du et al., 2008). Our GWAS did not detect signals at the *AOP2* loci, as GWAS models primarily detect marginal additive effects, so loci involved in epistatic interactions may leave little detectable signal when analyzed one variant at a time (Slim et al., 2020; Stamp et al., 2025; Zhou et al., 2013). Indeed, a GWAS of our key glucosinolate (2-OH-3-butenyl), which is known to be determined by epistasis among *AOP2*, *MAM1*, and *GS-OH* did not display a GWAS signal at *AOP2* (Katz et al. 2021).

All else equal, we might expect phenotypes (herbivore population growth here) with a simpler genetic architecture to evolve more rapidly than more complex ones determined by many genes (Macnair, 1991; Orr, 2000; Welch & Waxman, 2003). This is only true, however, when simple and complex phenotypes exhibit similar levels of standing genetic variation. Instead, we found that the key SNPs at these large-effect loci were all rare (< 3%), suggesting that there may be little standing genetic variation for selection to act on in natural populations of *A. thaliana*.

### Caveats

Although our study provides a rigorous analysis of the genetic architecture of a plant-herbivore interaction, there are several caveats to keep in mind. First, the controlled greenhouse conditions likely increased heritability estimates relative to field settings by reducing environmental variance; therefore, the estimates reported here should be interpreted as upper bounds (Weinig & Schmitt, 2004). Second, our assays of aphid population growth were measured during the vegetative rosette stage of *A. thaliana*, whereas the glucosinolate profiles we used from Katz *et al*. (2021) were measured in seeds. Importantly, the relative glucosinolate composition (chemotype) of *A. thaliana* is highly stable across tissues and developmental stages (Brown et al., 2003; Chan et al., 2010, 2011; Katz et al., 2021; Kliebenstein, Kroymann, et al., 2001; Kliebenstein, Lambrix, et al., 2001), but this could have introduced excess noise into our analysis. Third, we only examined the bottom-up effect of the plant on the herbivore, but not the top-down effect of the herbivore on the plant. Per-capita population growth of insect herbivores is often used as a metric of plant resistance (Kloth et al. 2012) and consumer-resource theory predicts that this bottom-up (per-capita) effect of the plant should be proportional to the top-down (per-capita) effect of the consumer (Murdoch et al. 2003). Understanding both the top-down and bottom-up effects is ultimately needed, however, to anticipate how eco-evolutionary dynamics will shape consumer-resource interactions. Finally, our genome-wide association analysis has implicated a number of novel candidate genes, but functional validation with reverse genetics (e.g., CRISPR-Cas9 knockouts, T-DNA insertion lines) is needed to move from association to causation (Bortesi & Fischer, 2015; Clauw et al., 2024; Lin et al., 2025; Ottaviani et al., 2025).

### Conclusion

Our work demonstrates that the strength of the plant-herbivore interaction between *A.thaliana and L. pseudobrassicae* is shaped by an oligogenic architecture. Interaction strength was influenced by epistasis among three plant defense genes (*AOP2*, *MAM1*, and *GS-OH*), which may simultaneously act to maintain genetic variation and constrain short-term evolutionary responses to selection (Barrett & Schluter, 2008). At the genome-wide level, a limited number of loci of additive effects contributed disproportionately to variation in herbivore response. However, genetic variation was limited at these large-effect loci, which would likely constrain their capacity to evolve in response to selection (Barrett and Schluter 2008). Together, these findings show that consumer-resource interactions can be shaped by additive and epistatic effects among a small number of loci, highlighting the need to integrate genomic effects into consumer-resource models to better anticipate their eco-evolutionary dynamics.

## Author Contributions

Following the CRediT model:

Conceptualization: CBPL, MB

Data Curation: CBPL, MB;

Formal Analysis: CBPL, FW, PEJ, AM, MB

Funding acquisition: MB;

Investigation: CBPL, EM, NA, SLL, MB

Methodology: CBPL, YS, MB;

Project Administration: CBPL

Resources: MB;

Supervision: MB

Validation: CBPL, SLL, MB

Visualization: CBPL, MB;

Writing-Original Draft: CBPL;

Writing - Review & Editing: CBPL, MB, YS, FW, PEJ, AM, EM, NA, SLL.

## Acknowledgements

We thank Mark Vellend and Peter Moffett for constructive feedback on a previous version; Centre SÈVE for acquiring the seeds from the 1001 genomes project; Frédérick St-Pierre for helping in the execution of the experiment; and Sylvain Lerat for facilitating the use of the greenhouse. We acknowledge the support of the Natural Sciences and Engineering Research Council of Canada (NSERC) Discovery Grant (MAB), Fonds de Recherche du Québec - Nature et Technologies (FRQNT) Relève Professorale Grant (https://doi.org/10.69777/342654) (MAB), Canadian Foundation for Innovation (CFI) John R. Evans Leaders Fund (MAB).

## Statement on inclusion

While this experimental research was conducted in a controlled greenhouse environment at the Université de Sherbrooke, Canada, we recognize the international context of the genetic resources used in this study. The plant accessions central to our work originated from diverse geographic regions, and we have cited the original sources and databases from which these materials were obtained to ensure appropriate attribution to the scientific communities that made them available. The research was primarily conducted within the bilingual (English and French) academic environment of Québec and was integrated into the local research community through the Complexe de recherche intégrative en sciences végétales et environnementales (CORSEVE) and through intellectual input from colleagues and trainees at the Université de Sherbrooke. Most authors are affiliated with the host Canadian institution, with one international co-author, Yasuhiro Sato, affiliated with Hokkaido University, Japan. We further recognize that the main campus of the Université de Sherbrooke is situated on the ancestral, unceded territory of the W8banaki Nation, the Ndakina, and Hokkaido University is situated on the ancestral land of Ainu People.

## Data availability

All data and code to reproduce the results reported in this manuscript will be made publicly available on GitHub and archived on Zenodo.

## Supporting Information

### Bulking seeds

Seeds were sown in 7 cm diameter pots filled with autoclaved Pro-Mix HP Mycorrhizae Potting soil and stratified in the dark at 4°C for two weeks. Plants were grown in growth chambers at 23°C with 16 hours light/8 hours dark for three weeks (thinned to one plant per pot one week after germination) before vernalization. Vernalization is a process in which plants require prolonged cold exposure to flower, and it was necessary for some of these diverse natural accessions (Laibach, 1943). To vernalize the seedlings, we grew them at 10°C with 8 hours of light and 4°C with 16 hours of dark, for 71 days to promote flowering. After vernalization, plants were isolated in individual sleeves to avoid cross-pollination and moved to the greenhouse for two months (22°C with 16 hours light/8 hours dark) to harvest their seeds following standard protocols (Rivero et al., 2014).

### Bulking aphids

To synchronize aphid age, we maintained a bulk colony of *L. pseudobrassicae* on radish (Raphanus sativus) host plants under controlled conditions. We began by planting radish seeds in 14 cm diameter pots filled with Pro-Mix Premium Potting Mix, using identical greenhouse conditions as for *A. thaliana* growth (22°C, 16h light/8h dark) to ensure environmental consistency. Three replicate systems were enclosed in BugDorm mesh cages (50 cm³) to prevent aphid dispersal. For nymph production, we transferred adult wingless aphids from our lab colonies to 2-week-old radish plants bagged in an organza mesh bag to prevent aphid dispersal (3 aphids/plant; 5 plants/pot; 5 pots/cage). We repeated this process every 10 days with fresh adult aphids from the same cohort to ensure all experimental nymphs came from adults of similar age. One day before adding the nymphs to the plants, adults were allowed to reproduce for 24 hours, during which each adult typically produced approximately 10 nymphs, yielding roughly 1000 first-instar nymphs per bulking cycle. We then collected first-instar nymphs using a fine brush and transferred 2 nymphs per *A. thaliana* plant.

### Follow-up experiment testing reproducibility of aphid mortality

To test whether genotype explains variation in the occurrence of zero aphid counts, we fitted binomial linear models with and without genotypes as a fixed effect on data from the main and follow-up experiments. Model fit was compared using likelihood ratio tests. In the main experiment (n=1,056 plants), 112 plants had zero aphids (10.6%). In the follow-up experiment, we included 13 *A. thaliana* accessions selected from the lower (n = 10 accessions) and upper (n = 3 accessions) tails of the primary aphid-count distribution, and we tested whether low aphid counts were reproducible under a second standardized assay. Across 158 plants, 7 plants (4.43%) had zero aphids. Adding genotype as a fixed effect, did not significantly improve model fit in either the main experiment (χ² = 353.3, df = 384, *p* = 0.883) or the follow-up experiment (χ² = 16.80, df = 12, *p* = 0.158), indicating that genotype does not explain variation in aphid absence in either experiment. While the zero-aphid rates differ between experiments (10.6% vs. 4.43%), both are relatively low, and neither shows evidence of genotype-specific effects. The lack of reproducible genotype-dependent patterns across independent experiments suggests that aphid mortality was due to handling during the experiment, as first-instar nymphs are delicate to manipulate.

---

**Figure S1.**
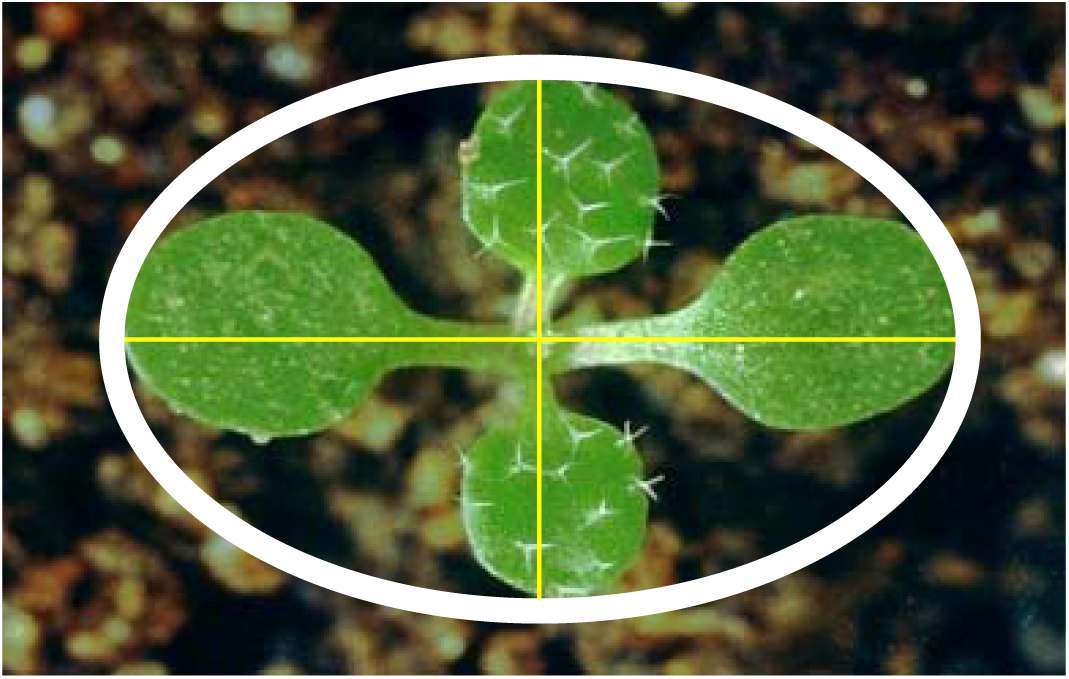
Measurement of plant size of an *Arabidopsis thaliana* seedling. The cotyledons (horizontal line) are almost round with a smooth surface. In contrast, the first true leaves (vertical line) are oval. As the plant grows, for some accessions, the leaves become more egg-shaped, and their surface appears rough due to trichomes. Modified from (Danish Agricultural Advisory Services, n.d.).

**Figure S2.**
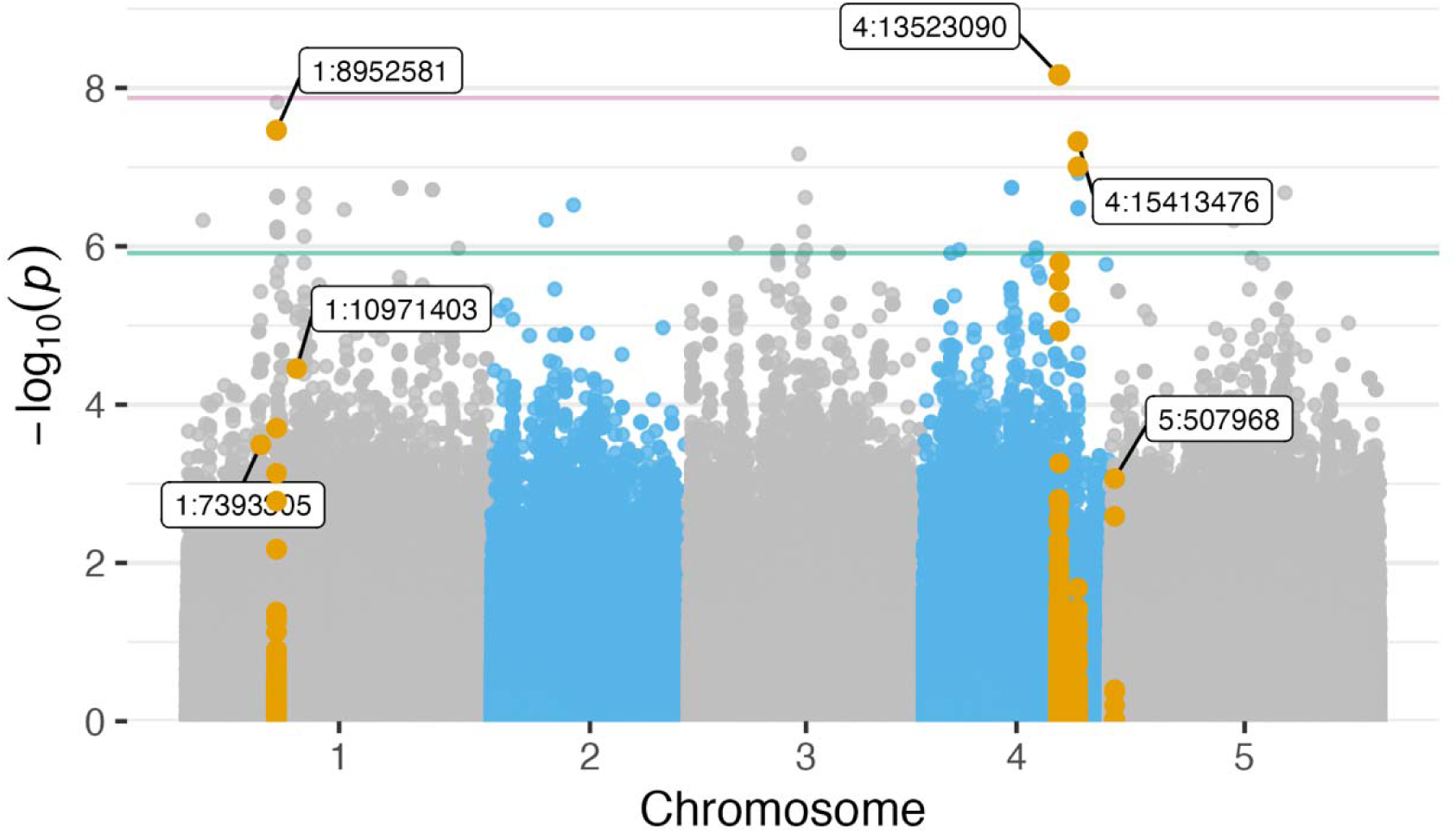
Genome-wide association study of herbivore per-capita population growth after adjusting for average plant size. Manhattan plot of single-nucleotide polymorphism (SNP) associations in *Arabidopsis thaliana* with herbivore per-capita population growth from a classic GWAS that controls for the average size of each accession. Each point represents a SNP, with orange points highlighting SNPs with posterior inclusion probabilities (PIP) > 0.01 (i.e., probability of having a non-zero effect), as inferred from a Bayesian sparse linear mixed model (BSLMM). Positions of SNPs with the lowest GWAS *p*-value in each of the six linkage disequilibrium blocks identified by BSLMM are labelled. The green and pink horizontal lines in (A) denote the genome-wide false-discovery rate and Bonferroni thresholds, respectively.

**Table S1.** Bayesian sparse linear mixed model (BSLMM) parameters for three independent MCMC chains (and their aggregated values) of herbivore per-capita growth rates on *Arabidopsis thaliana*.

| Statistic | Run | Mean | SD | Median | CI 2.5% | CI 97.5% |
| --- | --- | --- | --- | --- | --- | --- |
| PVE | Chain 1 | 0.506 | 0.137 | 0.502 | 0.249 | 0.782 |
|  | Chain 2 | 0.507 | 0.135 | 0.503 | 0.253 | 0.784 |
|  | Chain 3 | 0.502 | 0.139 | 0.499 | 0.239 | 0.782 |
|  | Aggregated | 0.505 | 0.137 | 0.501 | 0.247 | 0.783 |
| PGE | Chain 1 | 0.488 | 0.16 | 0.469 | 0.216 | 0.865 |
|  | Chain 2 | 0.488 | 0.158 | 0.469 | 0.222 | 0.863 |
|  | Chain 3 | 0.486 | 0.164 | 0.467 | 0.201 | 0.872 |
|  | Aggregated | 0.488 | 0.161 | 0.468 | 0.213 | 0.867 |
| $N_\gamma$ | Chain 1 | 5.286 | 3.997 | 4 | 2 | 13 |
|  | Chain 2 | 5.123 | 2.916 | 4 | 2 | 12 |
|  | Chain 3 | 7.103 | 14.668 | 4 | 2 | 26 |
|  | Aggregated | 5.837 | 8.983 | 4 | 2 | 13 |

**Table S2.**
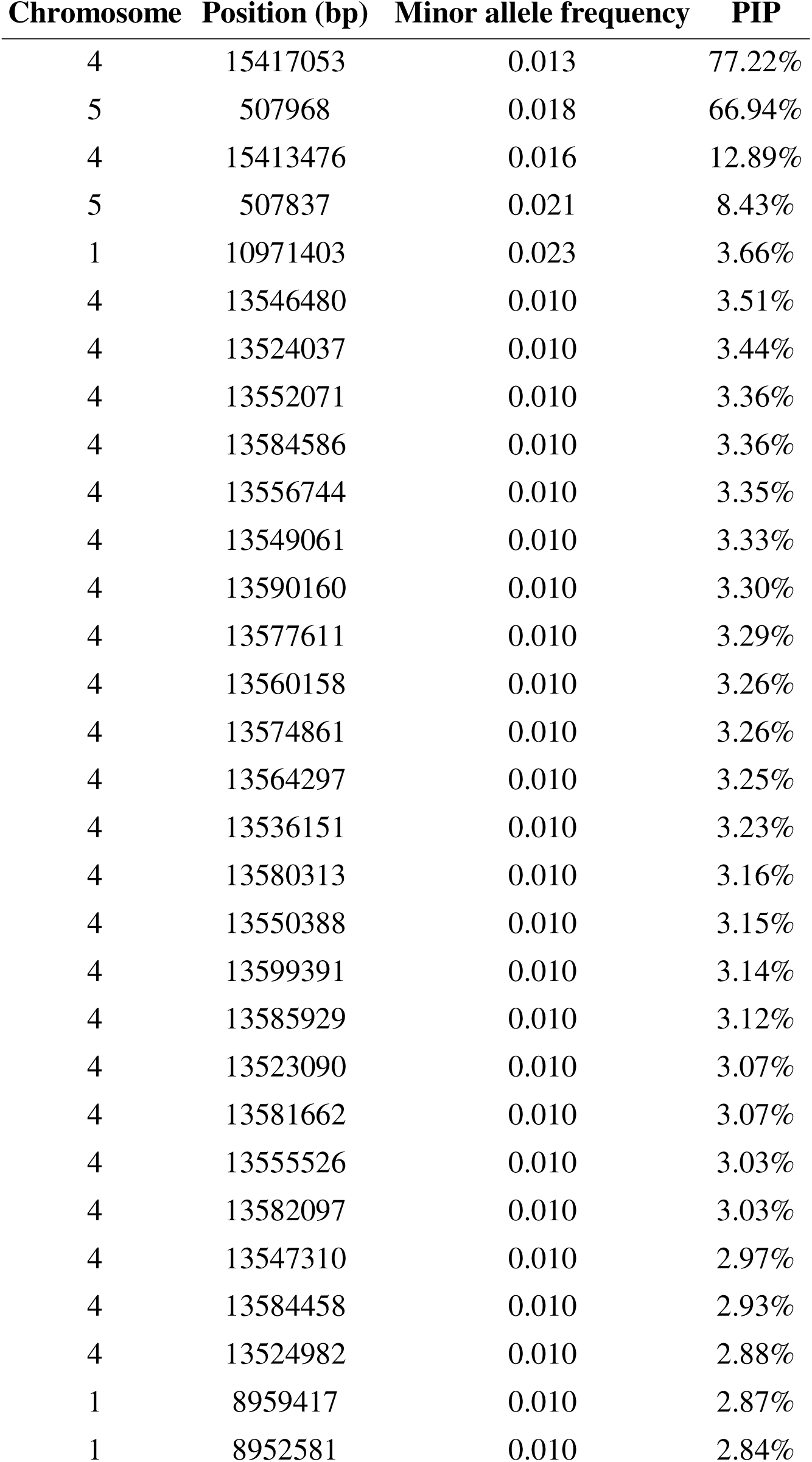
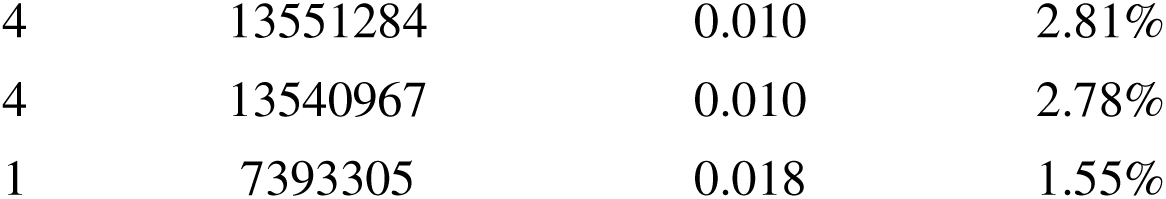
Single nucleotide polymorphisms (SNPs) identified by the Bayesian sparse linear mixed model with posterior inclusion probability (PIP) > 0.01. PIP quantifies the probability that a given SNP has a non-zero effect on herbivore population growth rates.

**Table S3.** Top 33 SNPs (PIP > 0.01) from BSLMM analysis of herbivore population growth rates in *A. thaliana* organized by linkage disequilibrium (LD) blocks.

| LD Block | Chr | Number of SNPs | Start position | End position | Range (kb) | ID of SNPs |
| --- | --- | --- | --- | --- | --- | --- |
| 1 | 1 | 1 | 7393305 | 7393305 | 0 | 1:7393305 |
| 2 | 1 | 2 | 8952581 | 8959417 | 6.84 | 1:8952581, 1:8959417 |
| 3 | 1 | 1 | 10971403 | 10971403 | 0 | 1:10971403 |
| 4 | 4 | 25 | 13523090 | 13599391 | 76.3 | 4:13523090,<br>4:13524037,<br>4:13524982,<br>4:13536151,<br>4:13540967,<br>4:13546480,<br>4:13547310,<br>4:13549061,<br>4:13550388,<br>4:13551284,<br>4:13552071,<br>4:13555526,<br>4:13556744,<br>4:13560158,<br>4:13564297,<br>4:13574861,<br>4:13577611,<br>4:13580313,<br>4:13581662,<br>4:13582097,<br>4:13584458,<br>4:13584586,<br>4:13585929,<br>4:13590160, 4:13599391 |
| 5 | 4 | 2 | 15413476 | 15417053 | 3.58 | 4:15413476, 4:15417053 |
| 6 | 5 | 2 | 507837 | 507968 | 0.131 | 5:507837, 5:507968 |

**Table S4.** Effects of *AOP3*-derived glucosinolates on aphid per-capita population growth rate. We followed the same analytical procedure as for the *AOP2*-derived compounds (see Methods).

| **Model** | **Predictor** | **Coefficient** | **95% CI** | ***p*-value** |
| --- | --- | --- | --- | --- |
| Total AOP3 | OHP3 + OHB4 | 0.023 | [-0.028, 0.074] | 0.37 |
| Specific AOP3 | OHP3 | 0.024 | [-0.027, 0.075] | 0.35 |
|  | OHB4 | -0.036 | [-0.159, 0.087] | 0.57 |

**Table S4.** Functional annotation of the 22 unique genes encompassed by linkage disequilibrium (LD) block 4 on chromosome 4 (positions 13,523,090 to 13,599,391 bp; 76.3 kb).

| Gene identifier | Function |
| --- | --- |
| AT4G26910 | Encodes the E2 subunit of the 2-oxoglutarate dehydrogenase; dihydrolipoamide succinyl transferase. |
| AT4G26920 | START (StAR-related lipid-transfer) lipid-binding domain-containing protein. |
| MYB97 | Encodes a putative transcription factor (MYB97); myb domain protein 97. |
| CAGE2 | Putative beta 1,3-galactosyltransferase that acts redundantly with CAGE1 during primary and secondary cell wall biosynthesis. |
| AT4G26950 | Protein of unknown function, DUF584; senescence regulator. |
| AT4G26960 | Hypothetical protein. |
| AT4G26965 | NADH: ubiquinone oxidoreductase, 17.2 kDa subunit. |
| ACO2 | Encodes an aconitase that catalyzes the conversion of citrate to isocitrate through a cis-aconitate intermediate; may participate in the TCA cycle and other primary metabolic pathways. |
| AT4G26980 | RNI-like superfamily protein. |
| AT4G26990 | Polyadenylate-binding protein interacting protein. |
| ATRBP45C | RNA-binding (RRM/RBD/RNP motifs) family protein. |
| EMB2788 | Ribosome 60S biogenesis amino-terminal protein; EMBRYO DEFECTIVE 2788. |
| AT4G27020 | Inositol-1,4,5-trisphosphate 5-phosphatase. |
| FADA | Encodes an unusual palmitate desaturase that introduces a delta-3 trans double bond at palmitate at the sn-2 position of phosphatidylglycerol. |
| VPS22 | EAP30/Vps36 family protein. |
| AT4G27050 | F-box/RNI-like superfamily protein. |
| TOR1 | Encodes a novel, plant-specific microtubule-associated protein that regulates cortical microtubule orientation and the direction of organ growth. |
| TSB2 | Tryptophan synthase beta-subunit 2; tryptophan synthase beta. |
| PDIL5-4 | Protein disulfide isomerase-like protein; member of the thioredoxin superfamily. |
| AT4G27090 | Cytoplasmic ribosomal protein L14; essential gene involved in gametophyte development and function. |
| AT4G27100 | Ypt/Rab-GAP domain of gyp1p superfamily protein. |
| COBL11 | GPI-anchored protein necessary for regulating pollen tube integrity; interacts with RALF-ANX-BUPS complex members and affects ROS levels and cell wall composition. |

## References

Alonso-Blanco, C., Andrade, J., Becker, C., Bemm, F., Bergelson, J., Borgwardt, K. M. M., Cao, J., Chae, E., Dezwaan, T. M. M., Ding, W., Ecker, J. R. R., Exposito-Alonso, M., Farlow, A., Fitz, J., Gan, X., Grimm, D. G. G., Hancock, A. M. M., Henz, S. R. R., Holm, S., … Zhou, X. (2016). 1,135 Genomes Reveal the Global Pattern of Polymorphism in Arabidopsis thaliana. Cell, 166(2), 481–491. 10.1016/J.CELL.2016.05.063/ATTACHMENT/D2E3D704-8EE7-4B34-B59C-81F2B4C47148/MMC2.PDF

Bailey, J. K., Wooley, S. C., Lindroth, R. L., & Whitham, T. G. (2006). Importance of species interactions to community heritability: a genetic basis to trophic-level interactions. Ecology Letters, 9, 78–85. 10.1111/j.1461-0248.2005.00844.x

Barbour, M. A., Fortuna, M. A., Bascompte, J., Nicholson, J. R., Julkunen-Tiitto, R., Jules, E. S., & Crutsinger, G. M. (2016). Genetic specificity of a plant-insect food web: Implications for linking genetic variation to network complexity. Proceedings of the National Academy of Sciences of the United States of America, 113(8), 2128–2133. 10.1073/PNAS.1513633113/SUPPL_FILE/PNAS.1513633113.SAPP.P DF

Barbour, M. A., Kliebenstein, D. J., & Bascompte, J. (2022). A keystone gene underlies the persistence of an experimental food web. Science, 376, 70–73. 10.1126/science.abf2232

Barbour, M. A., & Pérez-López, C. B. (2025). Linking plant genes to arthropod community dynamics: current progress and future challenges. Plant and Cell Physiology, 2025, 15. 10.1093/PCP/PCAF015

Barbour, M. A., Rodriguez-Cabal, M. A., Wu, E. T., Julkunen-Tiitto, R., Ritland, C. E., Miscampbell, A. E., Jules, E. S., & Crutsinger, G. M. (2015). Multiple plant traits shape the genetic basis of herbivore community assembly. Functional Ecology, 29(8), 995–1006. 10.1111/1365-2435.12409

Barrett, R. D. H., Rogers, S. M., & Schluter, D. (2008). Natural selection on a major armor gene in threespine stickleback. Science, 322(5899), 255–257. 10.1126/SCIENCE.1159978/SUPPL_FILE/BARRETT.SOM.PDF

Barrett, R. D. H., & Schluter, D. (2008). Adaptation from standing genetic variation. Trends in Ecology & Evolution, 23(1), 38–44. 10.1016/J.TREE.2007.09.008

Becks, L., Ellner, S. P., Jones, L. E., & Hairston Nelson G. J. G. (2010). Reduction of adaptive genetic diversity radically alters eco-evolutionary community dynamics. Ecology Letters, 13(8), 989–997. 10.1111/J.1461-0248.2010.01490.X

Benkman, C. W. (2013). Biotic interaction strength and the intensity of selection. Ecology Letters, 16(8), 1054–1060. 10.1111/ELE.12138

Berlow, E. L., Neutel, A. M., Cohen, J. E., De Ruiter, P. C., Ebenman, B., Emmerson, M., Fox, J. W., Jansen, V. A. A., Jones, J. I., Kokkoris, G. D., Logofet, D. O., Mckane, A. J., Montoya, J. M., & Petchey, O. (2004). Interaction strengths in food webs: issues and opportunities. Journal of Animal Ecology, 73(3), 585–598. 10.1111/J.0021-8790.2004.00833.X

Borg, M., Brownfield, L., Khatab, H., Sidorova, A., Lingaya, M., & Twella, D. (2011). The R2R3 MYB Transcription Factor DUO1 Activates a Male Germline-Specific Regulon Essential for Sperm Cell Differentiation in Arabidopsis. The Plant Cell, 23(2), 534–549. 10.1105/TPC.110.081059

Bortesi, L., & Fischer, R. (2015). The CRISPR/Cas9 system for plant genome editing and beyond. Biotechnology Advances, 33(1), 41–52. 10.1016/J.BIOTECHADV.2014.12.006

Brown, P. D., Tokuhisa, J. G., Reichelt, M., & Gershenzon, J. (2003). Variation of glucosinolate accumulation among different organs and developmental stages of Arabidopsis thaliana. Phytochemistry, 62(3), 471–481. 10.1016/S0031-9422(02)00549-6

Buniello, A., Macarthur, J. A. L., Cerezo, M., Harris, L. W., Hayhurst, J., Malangone, C., McMahon, A., Morales, J., Mountjoy, E., Sollis, E., Suveges, D., Vrousgou, O., Whetzel, P. L., Amode, R., Guillen, J. A., Riat, H. S., Trevanion, S. J., Hall, P., Junkins, H., … Parkinson, H. (2019). The NHGRI-EBI GWAS Catalog of published genome-wide association studies, targeted arrays and summary statistics 2019. Nucleic Acids Research, 47(D1), D1005–D1012. 10.1093/NAR/GKY1120

Burch, J., Chin, M., Fontenot, B. E., Mandal, S., McKnight, T. D., Demuth, J. P., & Blackmon, H. (2024). Wright was right: leveraging old data and new methods to illustrate the critical role of epistasis in genetics and evolution. Evolution, 78(4), 624– 634. 10.1093/EVOLUT/QPAE003

Bürkner, P. C. (2017). brms: An R package for Bayesian multilevel models using Stan. Journal of Statistical Software, 80. 10.18637/JSS.V080.I01

Chan, E. K. F., Rowe, H. C., Corwin, J. A., Joseph, B., & Kliebenstein, D. J. (2011). Combining Genome-Wide Association Mapping and Transcriptional Networks to Identify Novel Genes Controlling Glucosinolates in Arabidopsis thaliana. PLOS Biology, 9(8), e1001125. 10.1371/JOURNAL.PBIO.1001125

Chan, E. K. F., Rowe, H. C., & Kliebenstein, D. J. (2010). Understanding the evolution of defense metabolites in Arabidopsis thaliana using genome-wide association mapping. Genetics, 185(3), 991–1007. 10.1534/genetics.109.108522

Chang, C. C. C., Šlesak, I., Jordá, L., Sotnikov, A., Melzer, M., Miszalski, Z., Mullineaux, P. M., Parker, J. E., Karpińska, B., & Karpiñski, S. (2009). Arabidopsis chloroplastic glutathione peroxidases play a role in cross talk between photooxidative stress and immune responses. Plant Physiology, 150(2), 670–683. 10.1104/PP.109.135566

Clauw, P., Ellis, T. J., Liu, H.-J., & Sasaki, E. (2024). Beyond the Standard GWAS—A Guide for Plant Biologists. Plant and Cell Physiology. 10.1093/PCP/PCAE079

Cortez, M. H., Patel, S., & Schreiber, S. J. (2020). Destabilizing evolutionary and eco-evolutionary feedbacks drive empirical eco-evolutionary cycles. Proceedings of the Royal Society B: Biological Sciences, 287(1919). 10.1098/RSPB.2019.2298/85420

Daly, M. J., Rioux, J. D., Schaffner, S. F., Hudson, T. J., & Lander, E. S. (2001). High-resolution haplotype structure in the human genome. Nature Genetics 2001 29:2, 29(2), 229–232. 10.1038/ng1001-229

Davila Olivas, N. H., Kruijer, W., Gort, G., Wijnen, C. L., Van Loon, J. J. A., & Dicke, M. (2017). Genome-wide association analysis reveals distinct genetic architectures for single and combined stress responses in Arabidopsis thaliana. New Phytologist, 213, 838–851. 10.1111/nph.14165

Dong, S. S., He, W. M., Ji, J. J., Zhang, C., Guo, Y., & Yang, T. L. (2021). LDBlockShow: a fast and convenient tool for visualizing linkage disequilibrium and haplotype blocks based on variant call format files. Briefings in Bioinformatics, 22(4), 1–6. 10.1093/BIB/BBAA227

Du, J., Huang, Y. P., Xi, J., Cao, M. J., Ni, W. S., Chen, X., Zhu, J. K., Oliver, D. J., & Xiang, C. Bin. (2008). Functional gene-mining for salt-tolerance genes with the power of Arabidopsis. The Plant Journal : For Cell and Molecular Biology, 56(4), 653. 10.1111/J.1365-313X.2008.03602.X

Exposito-Alonso, M., Exposito-Alonso, M., Gómez Rodríguez, R., Barragán, C., Capovilla, G., Chae, E., Devos, J., Dogan, E. S., Friedemann, C., Gross, C., Lang, P., Lundberg, D., Middendorf, V., Kageyama, J., Karasov, T., Kersten, S., Petersen, S., Rabbani, L., Regalado, J., … Weigel, D. (2019). Natural selection on the Arabidopsis thaliana genome in present and future climates. Nature 2019 573:7772, 573(7772), 126–129. 10.1038/s41586-019-1520-9

Fujii, S., Tsuchimatsu, T., Kimura, Y., Ishida, S., Tangpranomkorn, S., Shimosato-Asano, H., Iwano, M., Furukawa, S., Itoyama, W., Wada, Y., Shimizu, K. K., & Takayama, S. (2019). A stigmatic gene confers interspecies incompatibility in the Brassicaceae. Nature Plants, 5(7), 731–741. 10.1038/S41477-019-0444-6

Galant, A., Preuss, M. L., Cameron, J. C., & Jez, J. M. (2011). Plant glutathione biosynthesis: Diversity in biochemical regulation and reaction products. Frontiers in Plant Science, 2(SEP). 10.3389/FPLS.2011.00045/ABSTRACT

Göbel, A. M., Zhou, S., Wang, Z., Tzourtzou, S., Himmelbach, A., Zheng, S., Pradillo, M., Liu, C., & Jiang, H. (2024). Mutations of PDS5 genes enhance TAD-like domain formation in Arabidopsis thaliana. Nature Communications 2024 15:1, 15(1), 9308-. 10.1038/s41467-024-53760-x

Goodey, N. A., Florance, H. V., Smirnoff, N., & Hodgson, D. J. (2015). Aphids Pick Their Poison: Selective Sequestration of Plant Chemicals Affects Host Plant Use in a Specialist Herbivore. Journal of Chemical Ecology, 41(10), 956–964. 10.1007/S10886-015-0634-2/FIGURES/4

Grafen, A. (1984). Natural Selection, Kin election and Group Selection. In J. R. Krebs & N. B. Davies (Eds.), Behavioural Ecology: An Evolutionary Approach (3). Blackwell Scientific Publications.

Groux, R., Stahl, E., Gouhier-Darimont, C., Kerdaffrec, E., Jimenez-Sandoval, P., Santiago, J., & Reymond, P. (2021). Arabidopsis natural variation in insect egg-induced cell death reveals a role for LECTIN RECEPTOR KINASE-I.1. Plant Physiology, 185(1), 240–255. 10.1093/PLPHYS/KIAA022

Guo, W.-J., Bundithya, W., & Goldsbrough, P. B. (2003). Characterization of the Arabidopsis metallothionein gene family: tissue-specific expression and induction during senescence and in response to copper. New Phytologist, 159. 10.1046/j.1469-8137.2003.00813.x

Hansen, B. G., Kerwin, R. E., Ober, J. A., Lambrix, V. M., Mitchell-Olds, T., Gershenzon, J., Halkier, B. A., & Kliebenstein, D. J. (2008). A Novel 2-Oxoacid-Dependent Dioxygenase Involved in the Formation of the Goiterogenic 2-Hydroxybut-3-enyl Glucosinolate and Generalist Insect Resistance in Arabidopsis. Plant Physiology, 148(4), 2096–2108. 10.1104/PP.108.129981

Harrison, X. A. (2014). Using observation-level random effects to model overdispersion in count data in ecology and evolution. PeerJ, 2(1). 10.7717/PEERJ.616

Hendry, A. P. (2013). Key questions in the genetics and genomics of eco-evolutionary dynamics. Heredity, 111(6), 456–466. 10.1038/hdy.2013.75

Hendry, A. P. (2017). Eco-evolutionary dynamics. Princenton University Press.

Holland, J. B., & Piepho, H.-P. (2024). Don’t BLUP Twice. G3: Genes, Genomes, Genetics, 14(12), 250. 10.1093/g3journal/jkae250

Hughes, M. A. (1991). The cyanogenic polymorphism in Trifolium repens L. (white clover). Heredity 1991 66:1, 66(1), 105–115. 10.1038/hdy.1991.13

Jie Lei, G., Yamaji, N., & Feng Ma, J. (2021). Two metallothionein genes highly expressed in rice nodes are involved in distribution of Zn to the grain. New Phytologist, 229, 1007–1020. 10.1111/nph.16860

Katz, E., Li, J. J., Jaegle, B., Ashkenazy, H., Abrahams, S. R., Bagaza, C., Holden, S., Pires, C. J., Angelovici, R., & Kliebenstein, D. J. (2021). Genetic variation, environment and demography intersect to shape Arabidopsis defense metabolite variation across Europe. ELife, 10. 10.7554/ELIFE.67784

Kawakatsu, T., Huang, S. C., Jupe, F., Sasaki, E., Schmitz, R. J. J., Urich, M. A. A., Castanon, R., Nery, J. R. R., Barragan, C., He, Y., Chen, H., Dubin, M., Lee, C. R., Wang, C., Bemm, F., Becker, C., O’Neil, R., O’Malley, R. C. C., Quarless, D. X. X., … Schork, N. J. (2016). Epigenomic Diversity in a Global Collection of Arabidopsis thaliana Accessions. Cell, 166(2), 492–505. 10.1016/J.CELL.2016.06.044

Kim, J. H., & Jander, G. (2007). Myzus persicae (green peach aphid) feeding on Arabidopsis induces the formation of a deterrent indole glucosinolate. The Plant Journal, 49(6), 1008–1019. 10.1111/J.1365-313X.2006.03019.X

Kinsella, R. J., Kähäri, A., Haider, S., Zamora, J., Proctor, G., Spudich, G., Almeida-King, J., Staines, D., Derwent, P., Kerhornou, A., Kersey, P., & Flicek, P. (2011). Ensembl BioMarts: a hub for data retrieval across taxonomic space. Database, 2011. 10.1093/DATABASE/BAR030

Kliebenstein, D. J. (2014). Quantitative Genetics and Genomics of Plant Resistance to Insects. Annual Plant Reviews Online, 47, 235–262. 10.1002/9781119312994.APR0511

Kliebenstein, D. J. (2017). Quantitative Genetics and Genomics of Plant Resistance to Insects. Annual Plant Reviews Online, 47, 235–262. 10.1002/9781119312994.APR0511

Kliebenstein, D. J., & Cacho, N. I. (2016). Nonlinear Selection and a Blend of Convergent, Divergent and Parallel Evolution Shapes Natural Variation in Glucosinolates. Advances in Botanical Research, 80, 31–55. 10.1016/BS.ABR.2016.06.002

Kliebenstein, D. J., Kroymann, J., Brown, P., Figuth, A., Pedersen, D., Gershenzon, J., & Mitchell-Olds, T. (2001). Genetic control of natural variation in Arabidopsis glucosinolate accumulation. Plant Physiology, 126(2), 811–825. 10.1104/PP.126.2.811

Kliebenstein, D. J., Lambrix, V. M., Reichelt, M., Gershenzon, J., & Mitchell-Olds, T. (2001). Gene duplication in the diversification of secondary metabolism: tandem 2-oxoglutarate-dependent dioxygenases control glucosinolate biosynthesis in Arabidopsis. The Plant Cell, 13(3), 681–693. 10.1105/TPC.13.3.681

Kloth, K. J., Thoen, M. P. M., Bouwmeester, H. J., Jongsma, M. A., & Dicke, M. (2012). Association mapping of plant resistance to insects. Trends in Plant Science, 17(5), 311–319. 10.1016/j.tplants.2012.01.002

Kroymann, J., Donnerhacke, S., Schnabelrauch, D., & Mitchell-Olds, T. (2003). Evolutionary dynamics of an Arabidopsis insect resistance quantitative trait locus. Proceedings of the National Academy of Sciences of the United States of America, 100(24), 14587–14592. 10.1073/PNAS.1734046100/SUPPL_FILE/4046FIG5.PDF

Kumar, S., Singh, Y. P., Singh, S. P., & Singh, R. (2017). Physical and biochemical aspects of host plant resistance to mustard aphid, Lipaphis erysimi (Kaltenbach) in rapeseed-mustard. Arthropod-Plant Interactions, 11(4), 551–559. 10.1007/S11829-016-9492-2/TABLES/2

Kwak, J. M., Moon, J. H., Murata, Y., Kuchitsu, K., Leonhardt, N., DeLong, A., & Schroeder, J. I. (2002). Disruption of a guard cell-expressed protein phosphatase 2A regulatory subunit, RCN1, confers abscisic acid insensitivity in Arabidopsis. The Plant Cell, 14(11), 2849–2861. 10.1105/TPC.003335

Lande, R., & Arnold, S. J. (1983). THE MEASUREMENT OF SELECTION ON CORRELATED CHARACTERS. Evolution, 37(6), 1210–1226. 10.1111/J.1558-5646.1983.TB00236.X

Lefcheck, J. S. (2015). PIECEWISESEM: Piecewise structural equation modelling in R for ecology, evolution, and systematics. 10.1111/2041-210X.12512

Lin, X. X., Gong, B. Q., Wang, F. Z., Wan, J. B., Xiong, X., & Li, J. F. (2025). Versatile Applications of CRISPR Based Programmable T DNA Integration in Plants. Plant Biotechnology Journal, 23(12), 5950. 10.1111/PBI.70353

Macnair, M. R. (1991). Why the evolution of resistance to anthropogenic toxins normally involves major gene changes: the limits to natural selection. Genetica, 84(3), 213–219. 10.1007/BF00127250/METRICS

McCann, K. S. (2012). Food Webs. Princeton University Press.

McPeek, M. A., & Peckarsky, B. L. (1998). LIFE HISTORIES AND THE STRENGTHS OF SPECIES INTERACTIONS: COMBINING MORTALITY, GROWTH, AND FECUNDITY EFFECTS. Ecology, 79, 867–879.

Murdoch, W. W., Briggs, C. J., & Nisbet, R. M. (2003). Consumer-Resource Dynamics (S. A. Levin & H. S. Horn, Eds.). Princenton University Press.

Nasser, J., Bergman, D. T., Fulco, C. P., Guckelberger, P., Doughty, B. R., Patwardhan, T. A., Jones, T. R., Nguyen, T. H., Ulirsch, J. C., Lekschas, F., Mualim, K., Natri, H. M., Weeks, E. M., Munson, G., Kane, M., Kang, H. Y., Cui, A., Ray, J. P., Eisenhaure, T. M., … Engreitz, J. M. (2021). Genome-wide enhancer maps link risk variants to disease genes. Nature 2021 593:7858, 593(7858), 238–243. 10.1038/s41586-021-03446-x

Nordborg, M., Borevitz, J. O., Bergelson, J., Berry, C. C., Chory, J., Hagenblad, J., Kreitman, M., Maloof, J. N., Noyes, T., Oefner, P. J., Stahl, E. A., & Weigel, D. (2002). The extent of linkage disequilibrium in Arabidopsis thaliana. Nature Genetics 2002 30:2, 30(2), 190–193. 10.1038/ng813

Orr, H. A. (2000). Adaptation and the cost of complexity. Evolution; International Journal of Organic Evolution, 54(1), 13–20. 10.1111/J.0014-3820.2000.TB00002.X

Orr, H. A. (2005). The genetic theory of adaptation: a brief history. Nature Reviews Genetics 2005 6:2, 6(2), 119–127. 10.1038/nrg1523

Ottaviani, L., Lefeuvre, R., Montes, E., Widiez, T., Giorni, P., Mithöfer, A., Marocco, A., & Lanubile, A. (2025). A loss-of-function of ZmWRKY125 induced by CRISPR/Cas9 improves resistance against Fusarium verticillioides in maize kernels. Plant Cell Reports, 44(7), 144-. 10.1007/S00299-025-03544-4/FIGURES/7

Patel, S., Cortez, M. H., & Schreiber, S. J. (2018). Partitioning the Effects of Eco-Evolutionary Feedbacks on Community Stability*. The American Naturalist, 191(3), 381–394. 10.1086/695834

Pradillo, M., Knoll, A., Oliver, C., Varas, J., Corredor, E., Puchta, H., & Santos, J. L. (2015). Involvement of the Cohesin Cofactor PDS5 (SPO76) During Meiosis and DNA Repair in Arabidopsis thaliana. Frontiers in Plant Science, 6(DEC). 10.3389/FPLS.2015.01034

Rausher, M. D., & Delph, L. F. (2015). Commentary: When does understanding phenotypic evolution require identification of the underlying genes? Evolution, 69(7), 1655–1664. 10.1111/EVO.12687

Rowe, H. C., & Kliebenstein, D. J. (2008). Complex Genetics Control Natural Variation in Arabidopsis thaliana Resistance to Botrytis cinerea. Genetics, 180(4), 2237–2250. 10.1534/GENETICS.108.091439

Sato, Y., Tezuka, A., Kashima, M., Deguchi, A., Shimizu-Inatsugi, R., Yamazaki, M., Shimizu, K. K., & Nagano, A. J. (2019). Transcriptional variation in glucosinolate biosynthetic genes and inducible responses to aphid herbivory on field-grown Arabidopsis thaliana. Frontiers in Genetics, 10(JUL), 469835. 10.3389/FGENE.2019.00787/BIBTEX

Seren, Ü., Grimm, D., Fitz, J., Weigel, D., Nordborg, M., Borgwardt, K., & Korte, A. (2017). AraPheno: a public database for Arabidopsis thaliana phenotypes. Nucleic Acids Research, 45(D1), D1054–D1059. 10.1093/nar/gkw986

Shipley, B. (2000). Cause and Correlation in Biology. Cause and Correlation in Biology. 10.1017/CBO9780511605949

Slim, L., Chatelain, C., Azencott, C. A., & Vert, J. P. (2020). Novel methods for epistasis detection in genome-wide association studies. PLOS ONE, 15(11), e0242927. 10.1371/JOURNAL.PONE.0242927

Speed, D., Cai, N., Johnson, M. R., Nejentsev, S., & Balding, D. J. (2017). Reevaluation of SNP heritability in complex human traits. Nature Genetics 2017 49:7, 49(7), 986–992. 10.1038/ng.3865

Stamp, J., Smith, S. P., Weinreich, D., & Crawford, L. (2025). Sparse modeling of interactions enables fast detection of genome-wide epistasis in biobank-scale studies. American Journal of Human Genetics, 112(9), 2198–2212. 10.1016/J.AJHG.2025.07.004

Stanton-Geddes, J., Yoder, J. B., Briskine, R., Young, N. D., & Tiffin, P. (2013). Estimating heritability using genomic data. Methods in Ecology and Evolution, 4(12), 1151–1158. 10.1111/2041-210X.12129

Steiner, C. C., Weber, J. N., & Hoekstra, H. E. (2007). Adaptive Variation in Beach Mice Produced by Two Interacting Pigmentation Genes. PLOS Biology, 5(9), e219. 10.1371/JOURNAL.PBIO.0050219

St-Pierre F., Ramírez I., & Barbour M. (2025). Empirically measuring eco-evolutionary stability. 10.22541/AU.176184008.86928529/V1

Taylor, A., Grapentine, S., Ichhpuniani, J., & Bakovic, M. (2021). Choline transporter-like proteins 1 and 2 are newly identified plasma membrane and mitochondrial ethanolamine transporters. Journal of Biological Chemistry, 296, 100604. 10.1016/j.jbc.2021.100604

Theologis, A., Ecker, J. R., Palm, C. J., Federspiel, N. A., Kaul, S., White, O., Alonso, J., Altafi, H., Araujo, R., Bowman, C. L., Brooks, S. Y., Buehler, E., Chan, A., Chao, Q., Chen, H., Cheuk, R. F., Chin, C. W., Chung, M. K., Conn, L., … Davis, R. W. (2000). Sequence and analysis of chromosome 1 of the plant Arabidopsis thaliana. Nature, 408(6814), 816–820. 10.1038/35048500

Thompson, J. N. (2005). The geographic mosaic of coevolution. University of Chicago Press.

Togninalli, M., Seren, Ü., Meng, D., Fitz, J., Nordborg, M., Weigel, D., Borgwardt, K., Korte, A., & Grimm, D. G. (2018). The AraGWAS Catalog: a curated and standardized Arabidopsis thaliana GWAS catalog. Nucleic Acids Research, 46(D1), D1150–D1156. 10.1093/NAR/GKX954

Uffelmann, E., Huang, Q. Q., Munung, N. S., de Vries, J., Okada, Y., Martin, A. R., Martin, H. C., Lappalainen, T., & Posthuma, D. (2021). Genome-wide association studies. Nature Reviews Methods Primers 2021 1:1, 1(1), 1–21. 10.1038/s43586-021-00056-9

Wang, H., Niu, Q. W., Wu, H. W., Liu, J., Ye, J., Yu, N., & Chua, N. H. (2015). Analysis of non-coding transcriptome in rice and maize uncovers roles of conserved lncRNAs associated with agriculture traits. The Plant Journal, 84(2), 404–416. 10.1111/TPJ.13018

Weinig, C., & Schmitt, J. (2004). Environmental Effects on the Expression of Quantitative Trait Loci and Implications for Phenotypic Evolution | BioScience | Oxford Academic. BioScience, 54(7), 627–365. https://academic.oup.com/bioscience/article-abstract/54/7/627/223523?redirectedFrom=fulltext

Welch, J. J., & Waxman, D. (2003). Modularity and the cost of complexity. Evolution; International Journal of Organic Evolution, 57(8), 1723–1734. 10.1111/J.0014-3820.2003.TB00581.X

Wootton, J. T., & Emmerson, M. (2005). Measurement of interaction strength in nature. Annual Review of Ecology, Evolution, and Systematics, 36(Volume 36, 2005), 419– 444. 10.1146/ANNUREV.ECOLSYS.36.091704.175535/CITE/REFWORK S

Yoshida, T., Jones, L. E., Ellner, S. P., Fussmann, G. F., & Hairston, N. G. (2003). Rapid evolution drives ecological dynamics in a predator–prey system. Nature 2003 424:6946, 424(6946), 303–306. 10.1038/nature01767

Zangerl, A. R., & Berenbaum, M. R. (2004). Genetic variation in primary metabolites of Pastinaca sativa; can herbivores act as selective agents? Journal of Chemical Ecology, 30(10), 1985–2002. 10.1023/B:JOEC.0000045590.28631.00/METRICS

Zhou, X., Carbonetto, P., & Stephens, M. (2013). Polygenic Modeling with Bayesian Sparse Linear Mixed Models. PLOS Genetics, 9(2), e1003264. 10.1371/JOURNAL.PGEN.1003264

Zhou, X., & Stephens, M. (2012). Genome-wide Efficient Mixed Model Analysis for Association Studies. Nature Genetics, 44(7), 821. 10.1038/NG.2310

